# Metabolic Interactive Nodular Network for Omics (MINNO): Refining and investigating metabolic networks based on empirical metabolomics data

**DOI:** 10.1101/2023.07.14.548964

**Authors:** Ayush Mandwal, Stephanie L. Bishop, Mildred Castellanos, Anika Westlund, George Chaconas, Ian Lewis, Jörn Davidsen

**Affiliations:** Department of Physics and Astronomy, University of Calgary, Calgary, AB, Canada; Department of Biological Sciences, University of Calgary, Calgary, AB, Canada; Department of Biochemistry and Molecular Biology, Cumming School of Medicine, Snyder Institute for Chronic Diseases, University of Calgary, Calgary, AB, Canada; Department of Microbiology, Immunology and Infectious Diseases, Cumming School of Medicine, Snyder Institute for Chronic Diseases, University of Calgary, Calgary, AB, Canada; Hotchkiss Brain Institute, University of Calgary, Calgary, AB, Canada

**Author notes:** To whom correspondence should be addressed. Correspondence may also be addressed to. Joint Authors (Ayush Mandwal, Stephanie L. Bishop).

## Abstract

Metabolomics is a powerful tool for uncovering biochemical diversity in a wide range of organisms, and metabolic network modeling is commonly used to frame results in the context of a broader homeostatic system. However, network modeling of poorly characterized, non-model organisms remains challenging due to gene homology mismatches. To address this challenge, we developed Metabolic Interactive Nodular Network for Omics (MINNO), a web-based mapping tool that takes in empirical metabolomics data to refine metabolic networks for both model and unusual organisms. MINNO allows users to create and modify interactive metabolic pathway visualizations for thousands of organisms, in both individual and multi-species contexts. Herein, we demonstrate an important application of MINNO in elucidating the metabolic networks of understudied species, such as those of the *Borrelia* genus, which cause Lyme disease and relapsing fever. Using a hybrid genomics-metabolomics modeling approach, we constructed species-specific metabolic networks for three *Borrelia* species. Using these empirically refined networks, we were able to metabolically differentiate these genetically similar species via their nucleotide and nicotinate metabolic pathways that cannot be predicted from genomic networks. These examples illustrate the use of metabolomics for the empirical refining of genetically constructed networks and show how MINNO can be used to study non-model organisms.

**GRAPHICAL ABSTRACT:** MINNO tool facilitates refining of metabolic networks, multi omics integration and investigation of cross-species interactions.

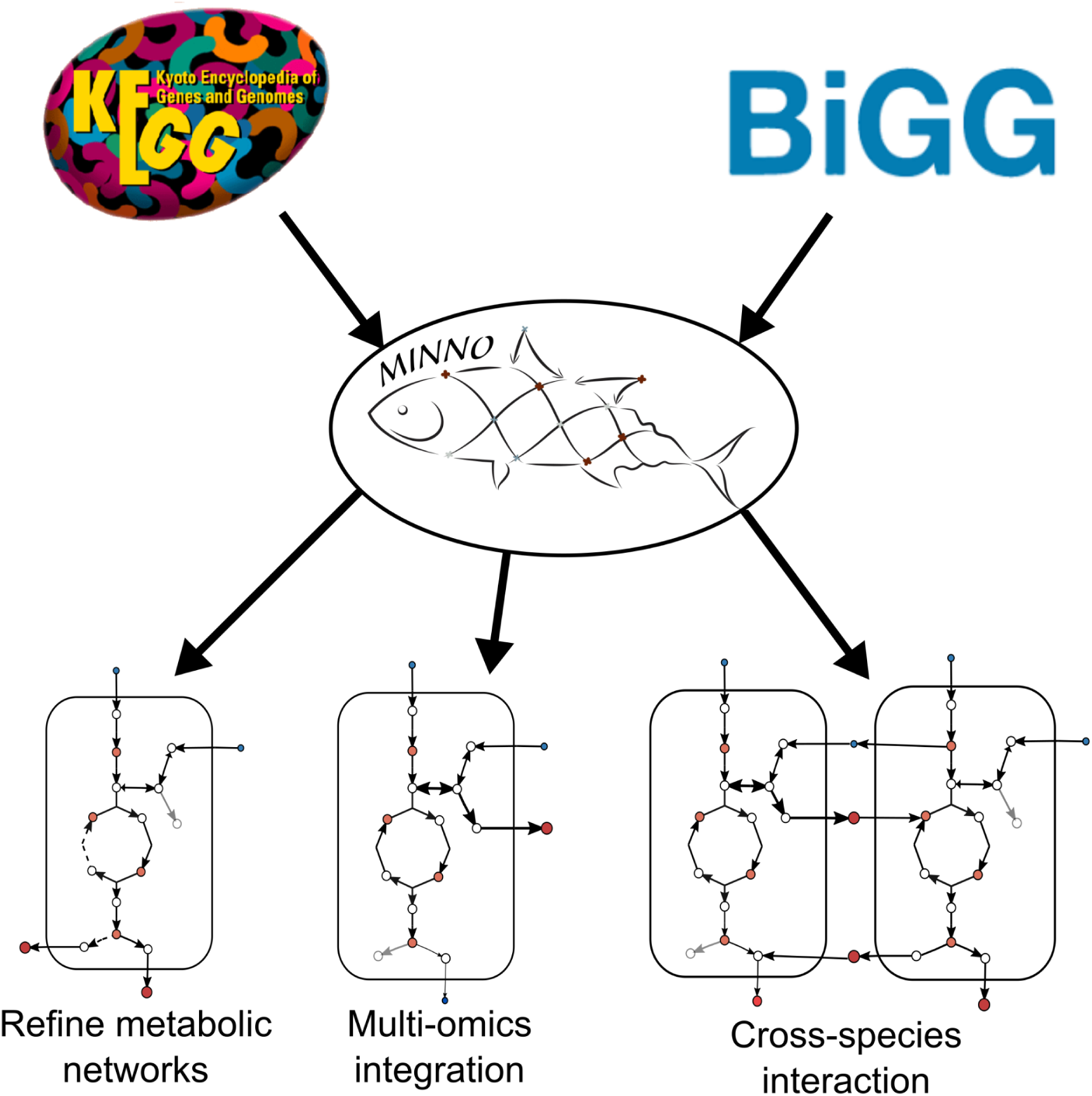

## INTRODUCTION

Metabolomics has emerged as a mainstream approach for investigating a diverse range of biological phenomena and exploring the molecular underpinnings of disease (1,2). One of the core underlying tools used in metabolomics research is metabolic network modeling, which is used to place metabolic data in the context of an organism’s overall metabolic network (3). Although the core elements of metabolism are shared between most living organisms (e.g., central carbon metabolism), species-to-species diversity can contribute to significant differences in nutritional preferences (4, 5). These differences are especially important in the context of evolution, where natural selection has driven organisms to streamline their metabolic networks according to the specific niches they inhabit (6).

Currently, most metabolic networks are derived from a handful of model organisms such as *Escherichia coli, Saccharomyces cerevisiae,* or *Mus musculus*. Species-specific networks are then constructed from genomic homology searches using tools such as the Prokaryotic Genome Annotation Pipeline (PGAP) (7,8), which identify the most likely enzyme for each reaction in the network (9). Although this strategy works well in species that are closely related to these model organisms, it is less effective when applied to species that are highly divergent from the original model (10). Mismatches due to poor homology result in missing enzymes in the metabolic network, which, without further data refinement, can be misinterpreted as metabolic deficiencies (11,12). This is a critical problem for understanding the evolution of microbes and for making inferences about the metabolic architecture of non-model organisms.

Another major challenge in investigating non-model organisms is that existing network visualization tools make it difficult to integrate multi-omics data to tune genomic networks. Although several visualization tools have been developed such as Escher (13), MetExploreViz (14), Omix (15), Cytoscape (16), CellDesigner (17), and PathVisio (18), they all suffer from a lack of scalability and reusability. None of these tools come with a generic base network architecture that can be used to build the network of any organism and they often require bioinformatics or coding skills to alter existing networks (15,17). When expanding metabolic networks by adding more pathways, users are typically required to manually add them one by one or reorganize the network if the initial layout is lacking (13,14,16,18).

To address these challenges, we developed the JavaScript-based web application Metabolic Interactive Nodular Network for Omics (MINNO). MINNO promotes network reusability by offering base networks that serve as a foundation for overlaying organism-specific networks. Moreover, it enables the integration of diverse metabolic pathways in a modular fashion, eliminating the need for coding or extensive reorganization of the entire combined network. These capabilities enhance the scalability of network construction and facilitate the empirical data-driven refinement of metabolomics networks. As a proof-of-concept, we used MINNO to conduct an empirical refinement of three metabolic pathways for three species of *Borrelia*, spirochetes that cause Lyme disease and relapsing fever in humans and other vertebrates (19, 20). *Borrelia* spirochetes follow a complex life cycle in which they are sequentially passed from ticks to a mammalian host (21, 22). These species are obligate parasites, and selective pressure has streamlined their metabolic networks to dispense with the biosynthesis of many metabolites that can be obtained directly from their host (19, 20). Using MINNO, we found evidence for metabolic streamlining and divergence among the Lyme disease and relapsing fever spirochetes resulting from these specific host/vector interactions.

In summary, the construction of metabolic networks for non-model organisms using genomics analysis is hindered by homology mismatches, which present a critical challenge in understanding microbial evolution and inferring their metabolic architecture. Existing visualization tools lack the necessary scalability and reusability features to effectively integrate multi-omics data into the network, thereby impeding network refinement. To address these limitations, MINNO utilizes a hybrid genomics-metabolomics strategy that incorporates metabolic boundary flux analysis, genomic network projection, and empirical refinement based on metabolic data and will thus facilitate the study of a significantly broader range of non-model species.

## MATERIAL AND METHODS

### Microbial growth and sample preparation

The *Borrelia* strains used in this study were *Borrelia burgdorferi* B31 5A4 (GCB921) (23), *Borrelia turicatae* 91E135 (GCB801), and *Borrelia parkeri* RML (GCB803) (24). For the extracellular metabolite temporal profiling dataset, all strains were propagated in BSK-II medium prepared in-house and supplemented with 6% rabbit serum (25). When the cultures approached log phase (1 to 5×10^7^), we diluted them in BSK-II with 6% rabbit serum to 1×10^5^ spirochetes/mL. We used 96-well plates (Life Science BRAND^TM^, Fisher Scientific, Toronto, ON, Canada) to grow triplicate samples of 250 μL for timepoints 0 and 72 hours and incubated them in a Forma^TM^ Series II Water-Jacketed CO_2_ Incubator (Thermo Scientific, Waltham, MA, USA) under conditions of 35°C and 1.5% CO_2_. At each timepoint, we transferred the cultures to 1.5 mL Eppendorf tubes and centrifuged them for 10 min at 13,000 rpm in a mini-centrifuge 5415 R (Eppendorf, Mississauga, ON, Canada), followed by removal of 100 μL of the supernatant and mixing it with an equal volume of 100% HPLC grade methanol (EMD Millipore, Oakville, ON, Canada). We then centrifuged the samples again under the above conditions and diluted the supernatant 1:10 with 50% LC-MS grade methanol (with 50% LC-MS grade water) to prepare the extracts for ultra-high-performance liquid chromatography-mass spectrometry (UHPLC-MS) analysis.

### Metabolomics analysis

We performed all metabolomics analyses at the Calgary Metabolomics Research Facility (CMRF), as described in detail in (26, 27). In brief, compounds were separated via hydrophilic interaction liquid chromatography (HILIC) with a 100 mm x 2.1 mm Syncronis™ HILIC LC column (2.1μm particle size; Thermo Fisher Scientific) using a Thermo Fisher Scientific Vanquish UHPLC platform and Thermo Scientific Q Exactive™ HF Hybrid Quadrupole-Orbitrap™ mass spectrometer. Raw MS data were converted to mzXML file format using MSConvert GUI software (28) and analyzed using El-Maven (El-MAVEN v0.12.0) (29). Metabolites were identified by comparison to an in-house metabolite library optimized for our instrumental setup (MetaSci, Toronto, ON, Canada) or quantified by comparison to standard solutions prepared from compounds ordered from Sigma-Aldrich or Acros Organics (now part of Thermo Fisher Scientific). Please refer to Table S1 for the CAS numbers of the compounds and select performance characteristics for the quantitative analysis.

For quantitative estimation of metabolic boundary fluxes (MBF), we used the following expression:

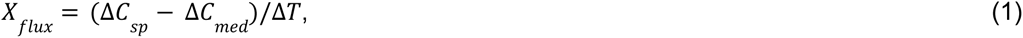

where ΔC_sp_ is the change in concentration of metabolite X in specific species samples while ΔC_med_ is the change in concentration of metabolite X in medium samples between t = 0hr and 72hr, such that ΔT=72hr. This expression ensures that the effect of medium evaporation and instrumental batch effects are normalized, limiting artefacts of the potential boundary flux profile. The significance of the quantified flux is estimated using the Student’s t-test with an α-level 0.05 for the measured change in the concentration for both species and medium samples.

## RESULTS and DISCUSSION

### Strategy: Network and data visualization using MINNO

The MINNO visualization tool facilitates both investigation and understanding of the complex interplay between genotypic and phenotypic features in omics data. MINNO is a JavaScript-based web application compatible with Google Chrome and Mozilla Firefox browsers. It uses the D3.js JavaScript library to create dynamic interactive visualizations in web browsers (30). The tool can load files, such as network files and data files, in JSON, XML, and CSV file formats, while it exports data in JSON, XML, PNG, and SVG formats for multiple applications. It has numerous built-in features that facilitate the creation of detailed network visualizations without the need to switch to multiple editing software tools. More details about MINNO can be found in the user manual that includes a tutorial developed for users to familiarize themselves with many of the tool’s features. MINNO is available open-source (under the MIT open-access license) at www.lewisresearchgroup.org/software.

MINNO comes with 66 base metabolic pathways from the KEGG database (31), covering all primary metabolic pathways that can be combined to build large-scale metabolic networks that include user-added reactions and features. Users can then superimpose an organism’s known metabolic pathway data from the KEGG database on these base metabolic pathways without the need to rebuild a network from scratch for each organism studied. The tool can also access metabolic network models from the Biochemical, Genetic and Genomic (BiGG) database (32). The tool accepts multi-omics data, such as metabolomics, proteomics, and fluxomics data, which can be integrated and visualized on the nodes and edges of the metabolic network. MINNO utilizes empirical data to facilitate the identification of missing reactions by providing users with the ability to investigate reactions pathway-by-pathway or by individual modules. The concept of modularity plays a crucial role in this process. Metabolic networks exhibit modularity as a network property, wherein a module or pathway consists of densely interconnected nodes compared to connections between different modules (33, 34). This modular structure enables the detection of missing reactions within metabolic networks by ensuring that nodes within each module are interconnected either with each other or with the surrounding environment. The concept of modularity is a fundamental aspect of metabolic networks and can be applied to metabolic networks of any species. However, except for a handful of model organisms, there are thousands of understudied species that have poorly constructed metabolic networks due to homology mismatch issues. The KEGG database currently includes over 8,794 species along with their respective metabolic pathways (31). By providing access to this extensive information, MINNO allows users to refine metabolic networks and explore individual species or interactions among multiple species.

### Network refinement strategy

In this example, we used MINNO for our metabolic network refinement strategy to understand metabolic differences among related microbial pathogens (Fig. 1). This strategy involves first culturing microbes in vitro and then sampling the cultures over specific time intervals so that metabolite intensities can be recorded as a function of time (Fig. 1A). Metabolites present in the cultures are then detected (in this example, by mass spectrometry) to generate temporal profiles of identified metabolites (Fig. 1B). The MINNO visualization tool takes metabolic base/ortholog network data (KGML files) from the KEGG database (Fig. 1C). Using the tool, users can superimpose an organism’s specific metabolic pathway onto base metabolic maps, which ultimately facilitates identification of potential missing reactions in the organism’s metabolic network as shown as grey edges. The user can then incorporate temporal metabolite intensity profiles and intra- or extracellular data onto the network to infer missing reactions by considering boundary fluxes and if available, the isotope labeling pattern, without resorting to complex mathematical modeling (Fig. 1D). In this figure, the dashed links represent the inferred missing reactions. Users can then search for genes and proteins corresponding to missing reactions using experimental or bioinformatics methods. The refined network with updated missing reactions can then be shared in KEGG format file (Fig. 1E). This approach allowed users to refine the networks of under-studied species such as *Borrelia* and find the necessary reactions to explain the metabolic profiles that were missing from their original annotated networks. MINNO can also be used for multi-omics data integration as it can incorporate gene, protein, metabolite, and flux data on the same metabolic network.

**Fig 1.**
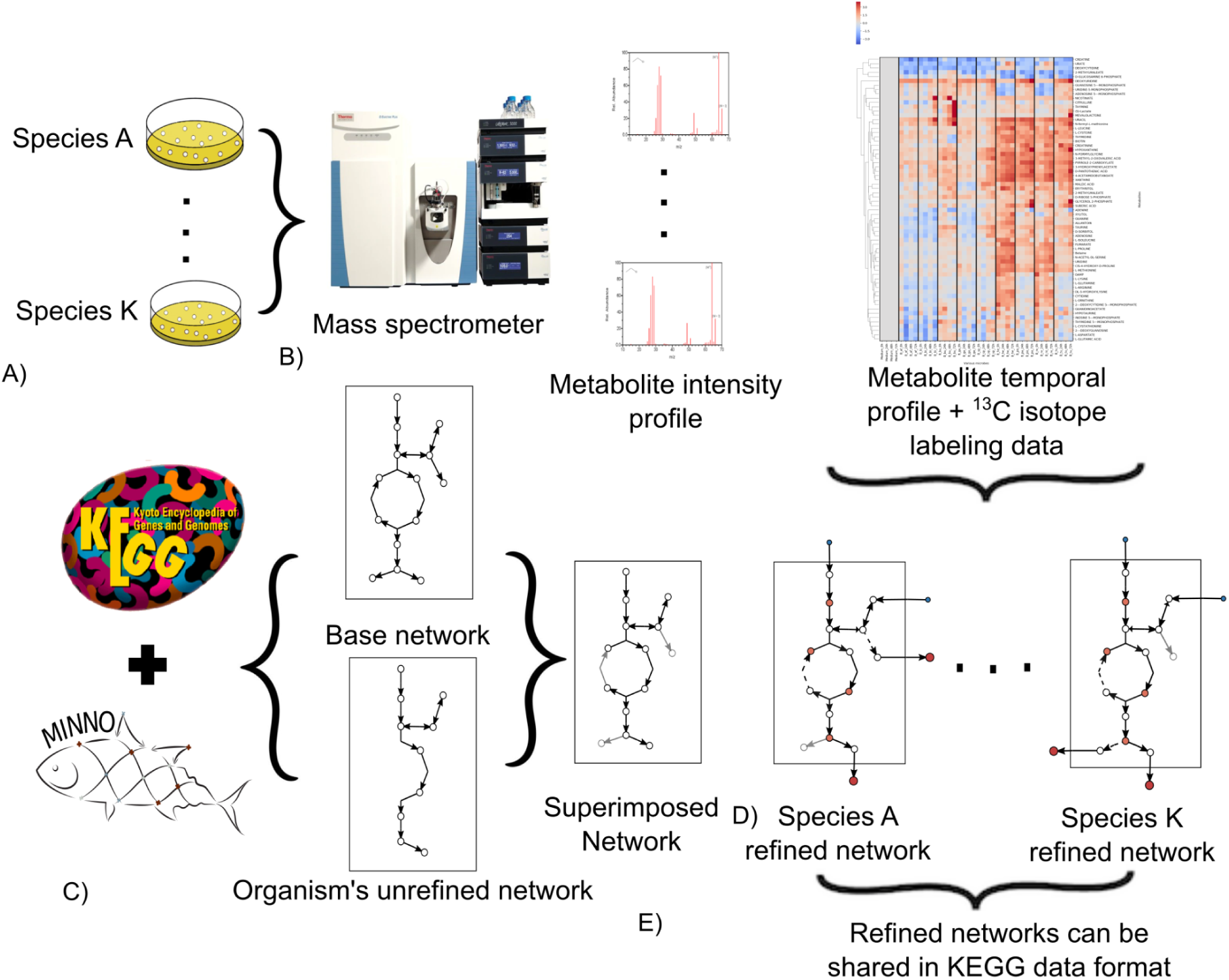
Schematic representation of the metabolic network refinement method, where microbes are grown in a specific growth medium to generate a temporal profile of metabolite intensities to refine the metabolic network of microbial species.

#### MINNO User interface

Fig. 2 highlights some key features of the web-based application MINNO. The built-in menu is located on the left side of the screen, where users can select base metabolic pathways or specific species pathways from the KEGG database and BiGG Models database. Metabolic pathways from the BiGG Models database do not have positional data. However, MINNO possesses the ability to self-organize the network using node repulsion and edge attractions from the D3.js JavaScript library (30), which is commonly employed in various other network visualization tools (14,16). Later, users can customize the network by dragging and aligning nodes in the network. Additionally, the tool allows users to superimpose base and organism-specific KEGG metabolic networks, which show annotated reactions as dark nodes and edges, while unannotated/missing reactions are depicted by light grey nodes and edges. This enables determining the potential missing reactions after experimental data is uploaded onto the network.

**Fig 2.**
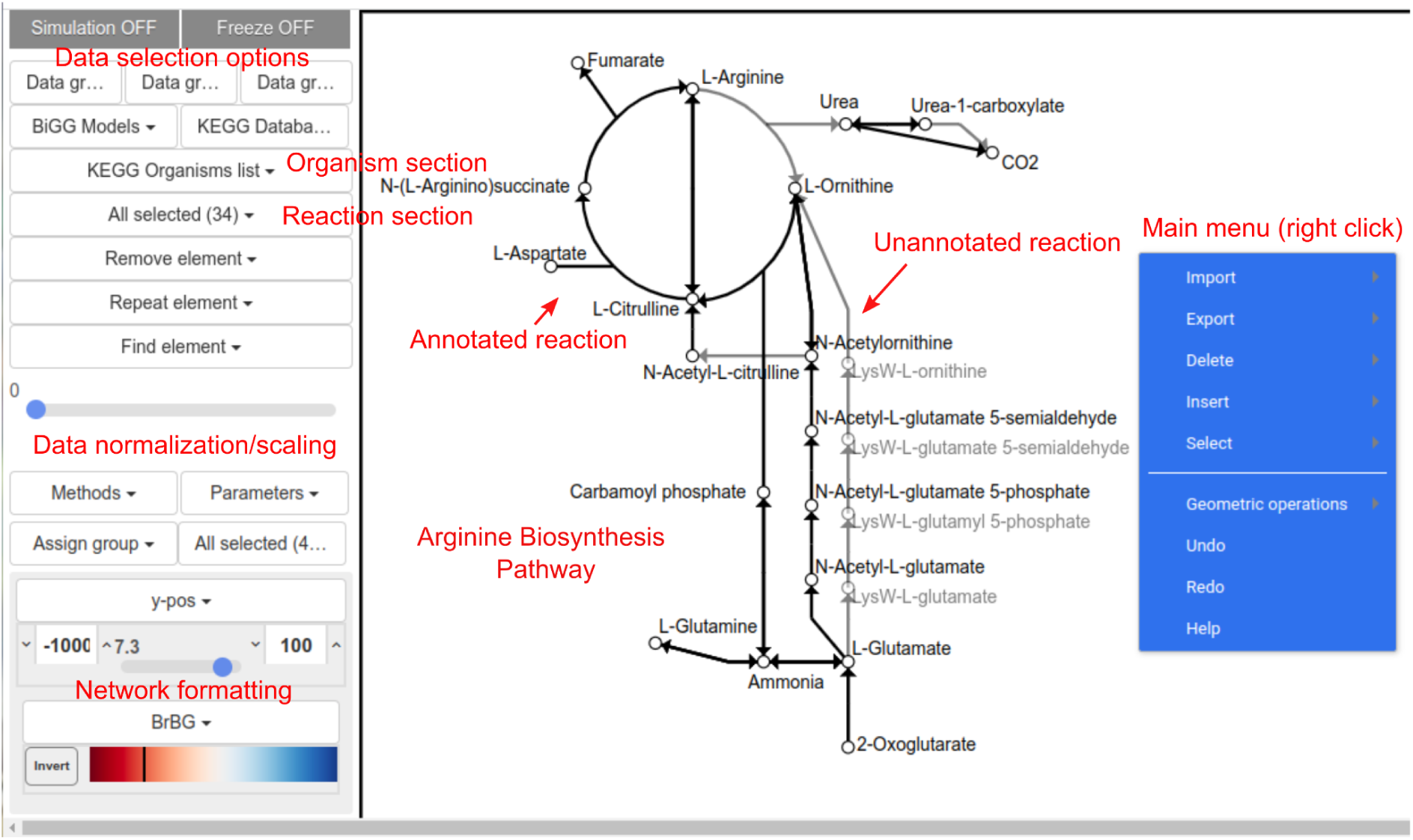
MINNO web-browser interface with key features highlighted in red.

### Refining nucleotide metabolic pathways using MINNO

We used MINNO to perform a metabolic network refinement analysis of three *Borrelia* species known to cause Lyme and relapsing fever diseases. By leveraging the modularity concept of metabolic networks and employing boundary flux analysis, we were able to identify missing reactions from the KEGG database for these species.

The purine metabolism of *B. burgdorferi* in the KEGG database is fragmented as depicted by solid links in Fig. 3A, as it lacks the classic purine salvage pathway (35). The consumption of both adenine and guanine by *B. burgdorferi* suggests the presence of purine transporters, which has recently been reported in the literature (36, 37, 38). By analyzing the boundary fluxes of cultured cells, we have identified missing reactions in purine metabolism from the KEGG database, indicated by dashed links in Fig. 3A. In contrast, the pyrimidine pathway for *B. burgdorferi* is relatively less fragmented in the KEGG database as shown by solid links in Fig. 3A. However, the boundary flux profile of this species suggests the presence of pyrimidine-nucleoside phosphorylase (*PnP*) based on the production of thymine from thymidine, as shown in Fig. 3A, which is missing in the KEGG database. Additionally, *B. burgdorferi* lacks ribonucleotide reductase, an enzyme responsible for converting ribonucleotides (for RNA synthesis) into deoxyribonucleotides (for DNA synthesis) (35). According to our data, *PnP* salvages deoxyribose sugars from thymidine for DNA synthesis. In summary, we used MINNO and empirical metabolomics data to identify eight reactions that are missing from the KEGG database. Subsequent publications have confirmed six of these missing purine reactions (Table 1). Furthermore, MINNO predicted four missing pyrimidine metabolism reactions from the KEGG database, all of which are supported by primary literature (Table 1).

**Fig. 3.**
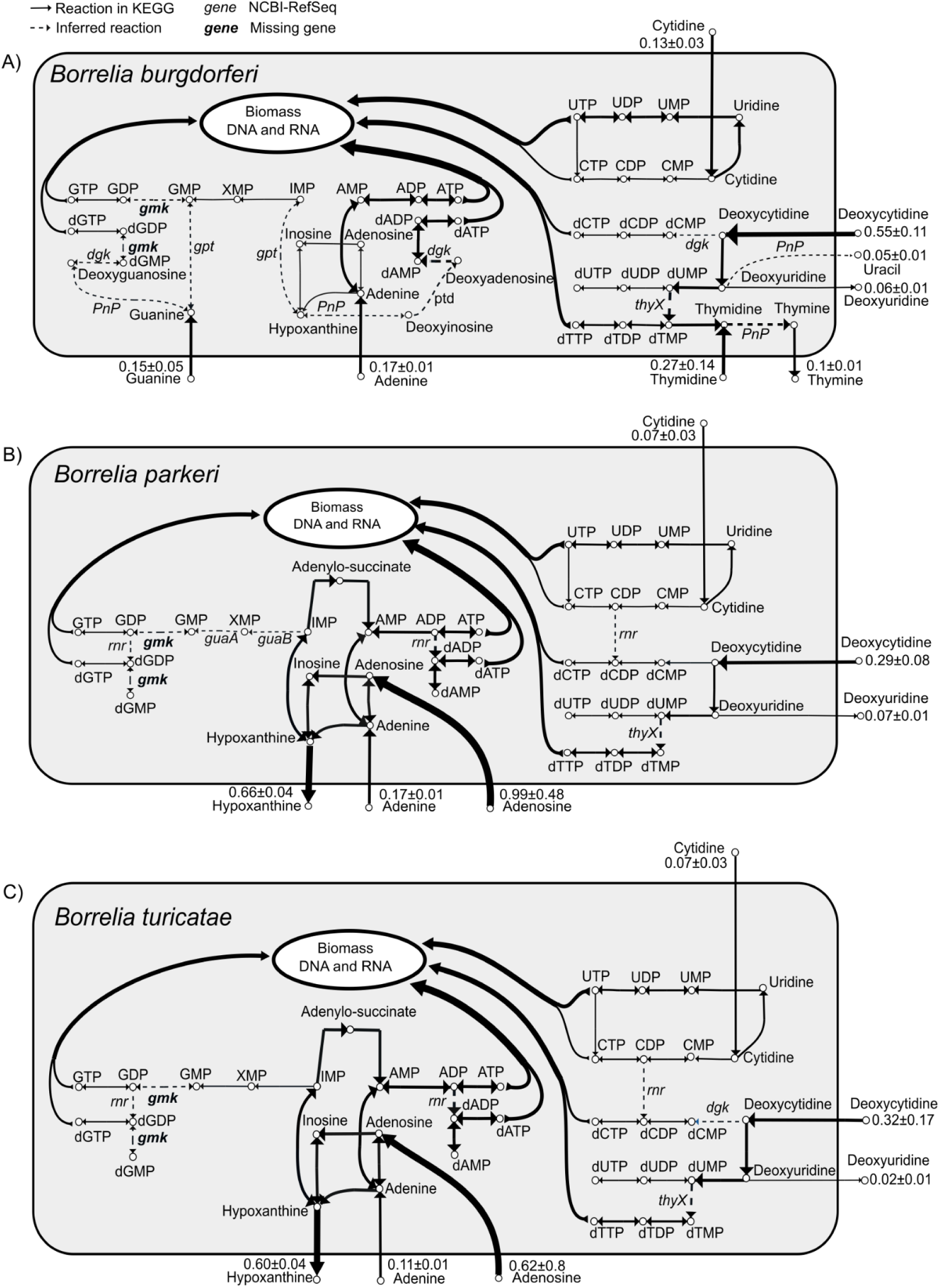
Purine and pyrimidine biosynthesis pathways of A) *B. burgdorferi*, B) *B. parkeri, and* C) *B. turicatae*. The solid black links represent annotated or documented reactions and dashed links are inferred missing reactions from KEGG database. The numerical values represent fluxes in μM/hr unit and the direction of flux is denoted by the arrow. The curved arrows highlight flow of metabolites toward the RNA and DNA biomass of various species. Thicker edges in the network implies higher flux and vice versa. All the genes were identified using the NCBI-RefSeq database and gene names are found in Table 1.

**Table 1.**
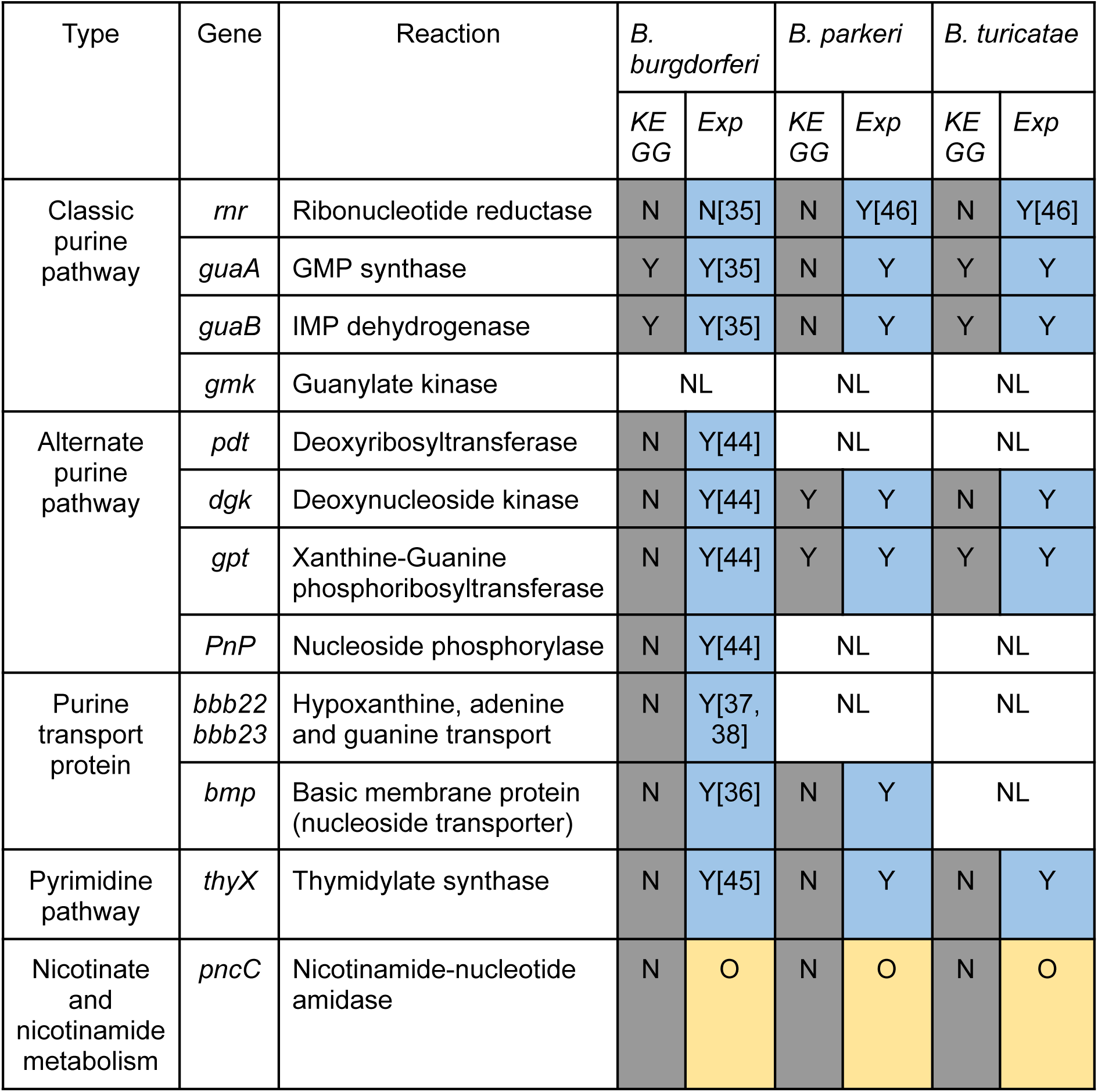
Tabulation of all reactions identified empirically using metabolomics data, including reactions that are documented in the KEGG database (grey) and experimentally validated (Exp; blue) with the corresponding reference. Y - reaction is present, N - reaction is absent, O - Orphan reaction and NL - no literature support exists. All identified reactions (blue) were also confirmed from the NCBI RefSeq database for all three species.

### Metabolic distinction between *Borrelia* species causing Lyme disease and relapsing fever

To better understand metabolic differences between *Borrelia* species, we focused on refining the metabolic networks of *Borrelia* species associated with relapsing fever: *B. parkeri* and *B. turicatae*. These species share genetic similarities, and as expected, their boundary flux profiles exhibit similarities as well (24) (Fig. 3 (B,C); Fig. 4). Similar to the purine metabolism of *B. burgdorferi,* purine metabolism of both *B. parkeri* and *B. turicatae* is fragmented as shown as solid links in Fig. 3 (B,C). However, unlike *B. burgdorferi*, both possess the classic purine salvage pathway. They both consume adenosine and adenine, and any excess purine is excreted as hypoxanthine. Interestingly, neither of the isolates studied has an annotated ribonucleotide reductase in the KEGG database. However, based on their boundary flux profiles, we anticipate that both *B. parkeri* and *B. turicatae* harbor a ribonucleotide reductase (*rnr*) (Fig. 3 B,C).

**Fig. 4.**
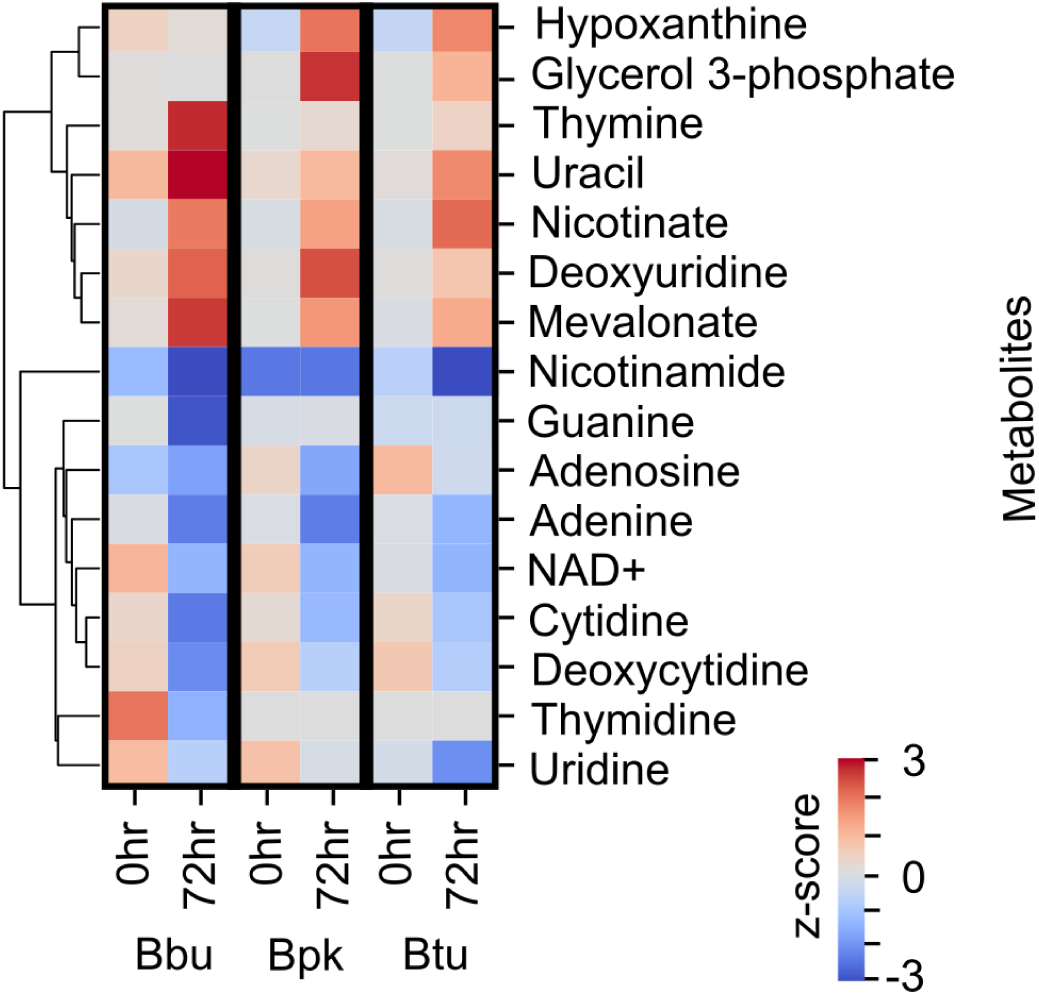
Heatmap showing the temporal profile of selected metabolite intensities across three *Borrelia* species bbu: *B. burgdorferi*, bpk: *B. parkeri* and btu: *B. turicatae* with respect to medium at 0hr and 72hr. The change in metabolite intensity across two different time-points (0 hr and 72 hr) has a *p*-value < 0.01 and the row z-score is shown through the color legend.

### Metabolic similarities between *Borrelia* species causing Lyme and Relapsing fever diseases

It is worth noting that these three *Borrelia* species also shared metabolic similarities. One common feature observed in all three species studied is the absence of the *thyX* gene in the KEGG database, as shown in Fig. 3. The *thyX* gene is essential in all three species for providing the necessary deoxyribonucleosides required for DNA synthesis. Another notable similarity is their deficiency in various biosynthetic pathways essential for the production of important vitamins and co-factors, such as nicotinate and nicotinamide. This deficiency highlights their reliance on salvaging precursors for NAD(P) synthesis from the host or surrounding environment. Our observations revealed that all three *Borrelia* species consume nicotinamide and NAD+ while excreting nicotinate out of the cells (Fig. 5). Notably, the net excretion of nicotinate exceeds the levels of nicotinamide consumed for each isolate. This suggests the possible presence of nicotinamide-nucleotide amidase (*pncC*). This salvage process also leads to the generation of essential molecules like PRPP and ATP, as well as the accumulation of potentially toxic substances such as ammonia.

**Fig. 5.**
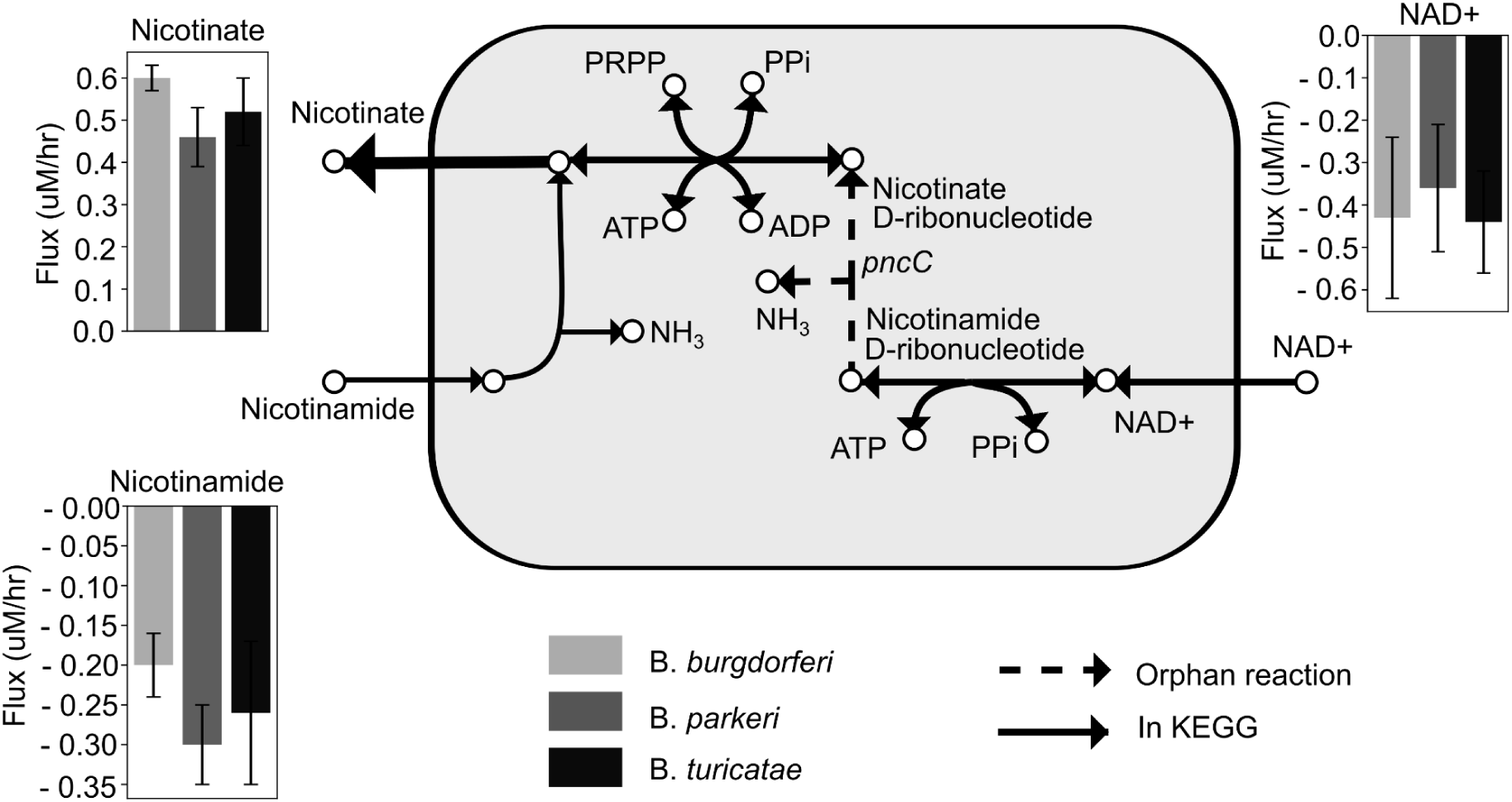
Refined nicotinate and nicotinamide metabolism. Solid links are annotated in the KEGG database while dashed links represent orphan reactions. Abbr. PRPP: Phosphoribosyl pyrophosphate, PPi: Diphosphate, NAD+: Nicotinamide adenine dinucleotide. The error bars represent standard deviation (sample size n=3).

In summary, we used MINNO to predict six reactions in purine metabolism for *B. parkeri* and *B. turicatae* that were missing from the KEGG database, with four of these predictions supported by primary literature (Table 1). MINNO was also used to identify two missing reactions in the KEGG database from the pyrimidine metabolism, although none of them are currently supported by primary literature. However, these predictions are supported based on homology matches through the PGAP pipeline from NCBI-RefSeq. For nicotinate metabolism, we predicted one reaction shared by all three *Borrelia* species, which is missing from the KEGG database (Table 1).

### Summary of functionality and applications

Overall, MINNO enables users to refine metabolic networks and integrate multi-omics data to provide a system level view of metabolic homeostasis. MINNO’s modular approach, whereby discrete metabolic pathway modules can be easily merged together, facilitates the creation of metabolic networks in diverse non-model organisms. It also allows users to visualize data on these merged metabolic pathways quickly and easily, without any coding required, facilitating a deeper understanding of complex multi-omics data in the context of the broader metabolic system. MINNO can support a variety of applications, such as FBA visualization to model more sophisticated genome-scale behaviors (39), mapping metabolic architecture in complex microbiome communities (40), investigating interspecies “cross-talking” interactions (41,42), and determining the molecular mechanisms of novel antibiotics (43).

## CONCLUSION

Here, we introduce MINNO, a new software tool that allows researchers to integrate genomic and empirical metabolomics data into a single software environment in order to build and refine metabolic networks. We illustrate the utility of this tool for identifying missing reactions within multiple metabolic pathways for *Borrelia* species. Using MINNO, we identified 18 missing reactions from the KEGG database, of which nine were supported by the primary literature. The remaining reactions show good homology as in the NCBI-RefSeq database (see Table 1). MINNO provides a tool that can be applied to any organism to systematically refine or investigate metabolic pathways. MINNO was designed to be inherently flexible for these diverse applications and support a wide range of input formats. We anticipate that it will be a useful asset for analyzing genome-wide knockouts, studying novel organisms that are divergent from typical model organisms, metabolic flux analysis, and visualization of metabolic networks.

## DATA AVAILABILITY

MINNO is available for free use (under the MIT license) on GitHub - https://lewisresearchgroup.github.io/MINNO/ as Lewis Research Group (LRG) software.

## AUTHOR CONTRIBUTIONS

Ayush Mandwal-Conceptualization, Formal analysis, Data curation, Software, Methodology, Validation, Writing—original draft.

Stephanie Bishop - Conceptualization, Validation, Methodology, Experimental, Writing—original draft.

Mildred Castellanos-Experimental

Anika Westlund - Validation

George Chaconas - Validation, Funding acquisition, Writing—review & editing

Ian Lewis-Conceptualization, Supervision, Validation, Funding acquisition, Writing—review & editing

Jörn Davidsen-Supervision, Funding acquisition, Writing—review & editing

## ACKNOWLEDGEMENTS

We thank Soren Wacker for providing necessary support in deploying the web application on GitHub for public use. IAL is supported by an Alberta Innovates Translational Health Chair and a grant from the Natural Sciences and Engineering Research Council of Canada (NSERC; DG04547). Metabolomics data were acquired at the Calgary Metabolomics Research Facility, which is supported by the International Microbiome Centre and the Canada Foundation for Innovation (CFI-JELF 34986).

